# ViDGER: An R package for integrative interpretation of differential gene expression results of RNA-seq data

**DOI:** 10.1101/268896

**Authors:** Adam McDermaid, Brandon Monier, Jing Zhao, Qin Ma

## Abstract

Differential gene expression (**DGE**) is one of the most common applications of RNA-sequencing (RNA-seq) data. This process allows for the elucidation of differentially expressed genes (**DEGs**) across two or more conditions. Interpretation of the DGE results can be non-intuitive and time consuming due to the variety of formats based on the tool of choice and the numerous pieces of information provided in these results files. Here we present an R package, **ViDGER** (Visualization of Differential Gene Expression Results using R), which contains nine functions that generate information-rich visualizations for the interpretation of DGE results from three widely-used tools, *Cuffdiff*, *DESeq2*, and *edgeR*.

## Introduction

Next-generation sequencing techniques enable researchers to access far more massive amounts of data than previously available. Specifically, RNA-seq procedures provide a plethora of information regarding the genetic expression levels of various organisms across multiple conditions at a high resolution [1, 2]. Naturally arising from this information is the concept of DEGs, which are genes that have expression levels determined to be significantly differentially expressed across two or more conditions. *Cuffdiff* [3, 4], *edgeR* [5], and *DESeq2* [6] are three widely-used tools to determine which genes are differentially expressed, based on quantifications of expressed genes derived from computational analyses of raw RNA-seq reads (e.g., mapping [7–15] and assembly [16–21]). Each of the three has been shown to be among the highest performing tools for DGE analysis of RNA-seq data [22–24] and contribute to the highest number of citations for DGE tools (Table 1), representing roughly 80% of all cited DGE tools. However, interpreting the results files from each program is not entirely intuitive, especially for researchers who have limited computational backgrounds.

One of the best ways to provide a summary of the DGE results is to generate figures, giving a global representation of the expression changes across multiple conditions. The three tools create output files sharing some information, such as mean gene expression across replicates for each sample, *log*_2_ fold change (*lfc*), and adjusted *p*-value. However, these output files have many differences in content and structure, which makes generating comprehensive visualizations time-intensive and potentially challenging task. *CummeRbund* [25] is an available tool to generate visualizations for *Cuffdiff* outputs but has no functionality for users of *edgeR* and *DESeq2*. This limited functionality leaves many researchers with no readily available method to create visualizations for their DGE results. To remediate this issue, we have created an R package, **ViDGER**, to assist users in generating publication-quality visualizations from *Cuffdiff*, *edgeR*, and *DESeq2* capable of providing valuable insight into their generated DGE results.

**Table 1.**
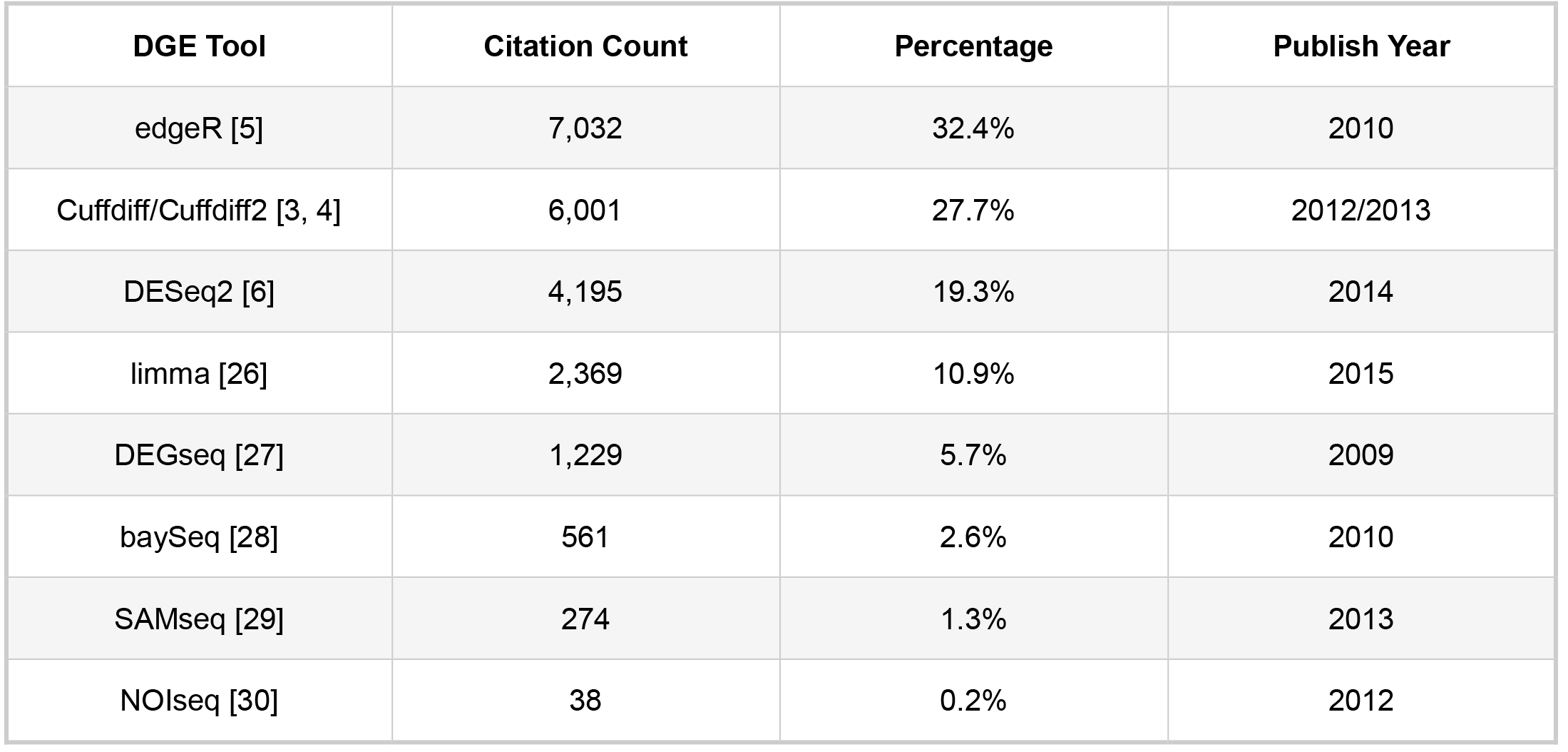
Citation counts, percentages of commonly referenced DGE tool citations, and year of release for edgeR [5], Cuffdiff/Cuffdiff2 [3, 4], DESeq2 [6], limma [26], DEGseq [27], baySeq [28], SAMseq [29], and NOIseq [30]. All counts were tabulated using the Google Scholar citation counts for the respective tool references as of Feb. 2, 2018.

| DGE Tool | Citation Count | Percentage | Publish Year |
| --- | --- | --- | --- |
| *edgeR* [5] | 7,032 | 32.4% | 2010 |
| *Cuffdiff*/*Cuffdiff2* [3, 4] | 6,001 | 27.7% | 2012/2013 |
| *DESeq2* [6] | 4,195 | 19.3% | 2014 |
| *limma* [26] | 2,369 | 10.9% | 2015 |
| *DEGseq* [27] | 1,229 | 5.7% | 2009 |
| *baySeq* [28] | 561 | 2.6% | 2010 |
| *SAMseq* [29] | 274 | 1.3% | 2013 |
| *NOIseq* [30] | 38 | 0.2% | 2012 |

This package can generate six different types of expression-based visualizations—boxplots, scatterplots, DEG counts, MA plots, volcano plots, and Four-way plots—as shown in Figure 1 and Examples S1-S9. Additionally, matrices of all pair-wise comparisons can be generated with scatterplots, MA plots, and volcanoplots (Examples S4, S7, and S9, respectively). All the visualizations can be classified into two tiers, with the Tier 1 functions (Figure 1A-C, Examples S2-S5) representing more basic information, whereas the Tier 2 functions (Figure 1D-F, Examples S6-S10) being used to derive more advanced information with *p*-values, fold changes, and mean expression values (Method S1). All generated figures and extracted data can then be saved and used for further purposes, including reports and publications.

**Figure. 1.**
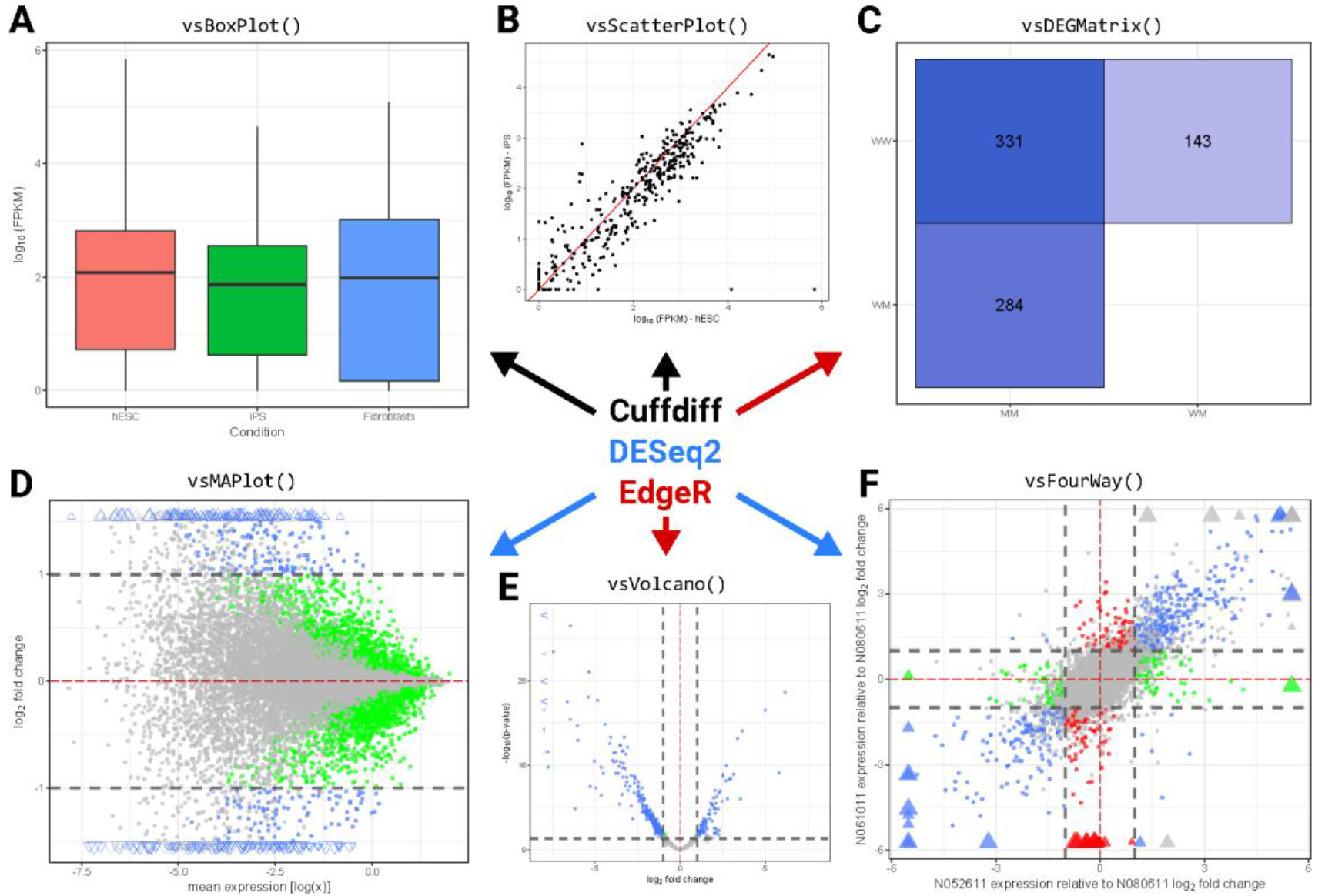
(A) Boxplot generation of RNA-seq data using *vsBoxplot*; (B) scatterplot generation using *vsScatterPlot*; (C) differential gene expression matrix using *vsDEGMatrix*; (D) MA plot generation using *vsMAPlot*; (E) volcano plot generation using *vsVolcano*; (F) four-way plot generation using *vsFourWay*. Arrow and text color refer to visualizations generated using *Cuffdiff* data (black), *DESeq2* data (blue), and *edgeR* data (red).

## Functions and Methods

Nine functions are included in **ViDGER**, each of which is capable of using *Cuffdiff*, *DESeq2*, and *edgeR* objects. Included in the ViDGER package are three toy datasets representing the three DGE tools object types. Specifically, *df.cuff* is based on *Cuffdiff* data from the *cummeRbund* package [25]; *df.deseq* is a *DESeqDataSet* object based on gene expression data from the *pasilla* package [31]; *df.edger* is an example *DGEList* object derived from the *edgeR* package (Example S1). In addition to the toy data sets, we tested ViDGER on five real-world data sets, consisting of one human, one *M. domestica*, and three *V. riparia* datasets (Example S1). It is important to note that the input data for this package should be the direct output and of one of the classes corresponding to the specific tool used (DESeqDataSet, DGEList or other edgeR objects, or Cuffdiff object) and not a basic matrix or data frame containing the results of these tools. The following examples are illustrated using the *df.deseq* object, with full demonstrations with the *Cuffdiff, DESeq2, and edgeR* objects found in the supplementary file (Examples S2-S10).

### Tier I Functions

**(i) *vsBoxPlot*** visualizes *log*_10_ distributions for treatments in an experiment as box and whisker diagrams (Figure 2, Example S2), where only the data frame and analytical type are needed unless using a DESeq2 object where the factor is also required. This figure is useful for determining the distribution of mapped read counts for each treatment in an experiment and can highlight specific samples that have distributions differing significantly from what is expected or what is displayed with the other samples. Visualizing this information can provide insight into the base quality of the read distributions to ensure semi-consistent sample-based quality levels. The *DESeq2* object (*df.deseq*) is used in the following example, and the factor variable, *d.factor*, for the treatments need to be specified.

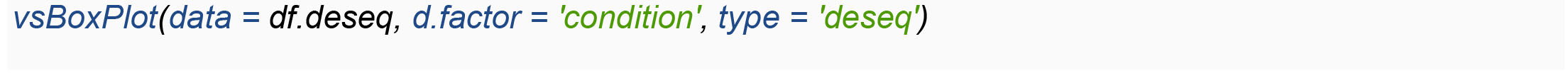

**Figure 2.**
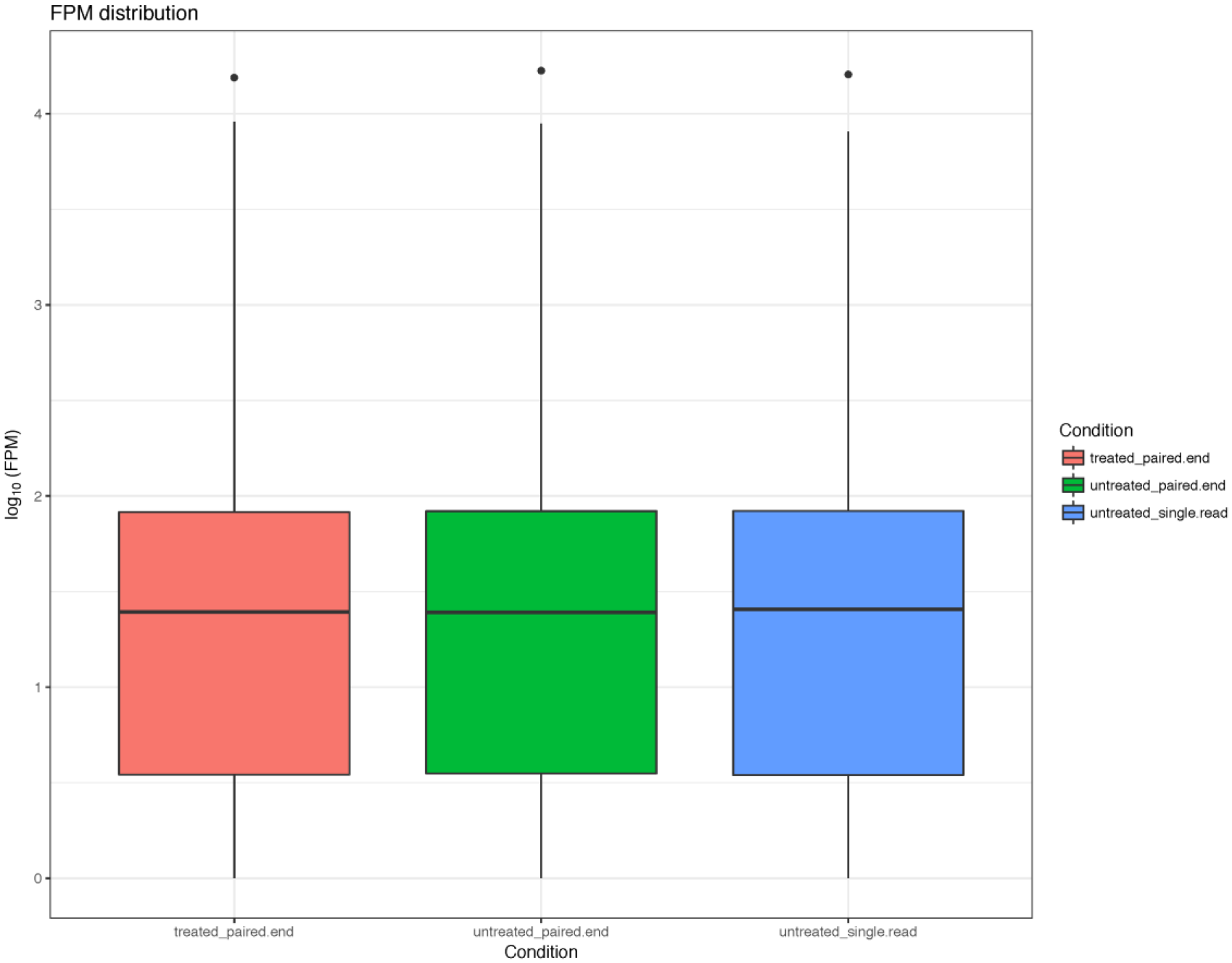
Visualization generated by the *vsBoxPlot* function from the ViDGER package using a DESeq2 dataset, requiring a dataset, factor type, and appropriate tool type. Optional parameters include inclusion/exclusion of the main title, legend, and grid.

**(ii) *vsScatterPlot*** creates a scatterplot of *log*_10_ comparison of either FPKM (Reads Per Kilobase of transcript per Million mapped reads) or CPM (cost per thousand impressions) measurements for two treatments depending on analytical type (Figure 3, Example S3). This function can be used to compare measurements of mapped reads to transcripts from two treatments, which allows for a global view of the expression similarity between the two selected treatments. Scatterplots that generate most data points falling along the diagonal indicate more similar expression patterns for the two treatments, whereas data points falling further from the diagonal would indicate relatively less similar expression levels. By stating *x* and *y* treatment variables and/or the data source, we can generate a scatterplot of the pairwise *x* vs. *y* comparison.

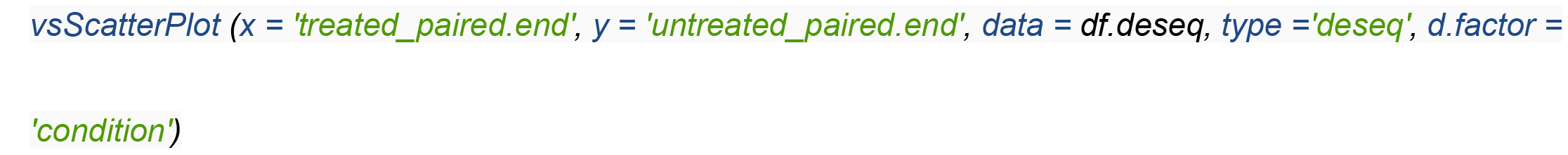

**Figure 3.**
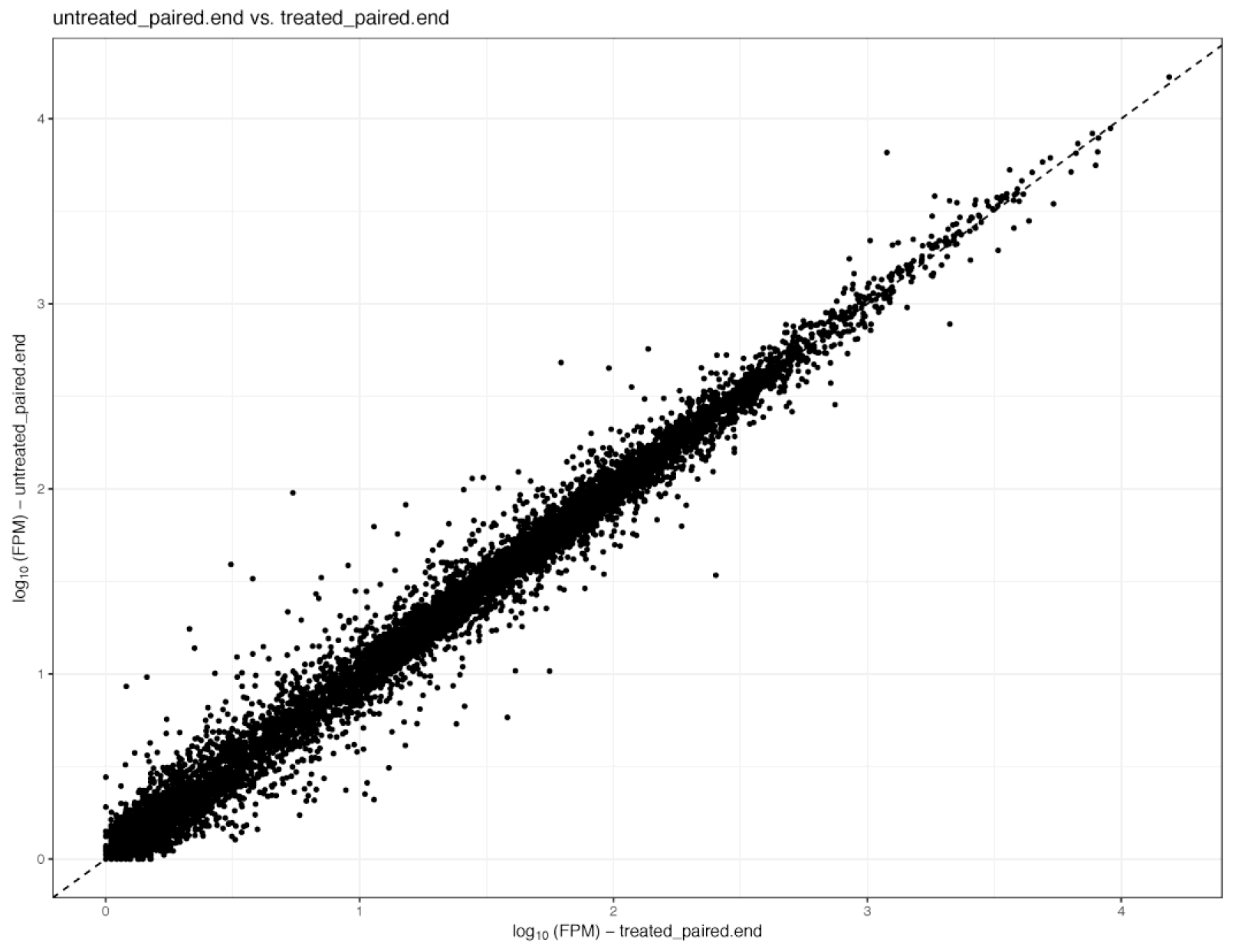
Visualization generated by the *vsScatterPlot* function from the ViDGER package using a DESeq2 dataset, requiring a dataset, factor type, two factor levels, and appropriate tool type. Optional parameters include inclusion/exclusion of the main title and grid.

**(iii) *vsScatterMatrix*** generates a matrix of scatterplots for all possible treatment combinations with additional distribution information (Figure 4, Example S4). In addition to the scatterplots which are generated as with the *vsScatterPlot* function, the matrix option provides FPKM/CPM distributions for each sample and correlation values for each pairwise comparison. This approach allows for a view of each relative expression pattern and correlation all in one visualization.

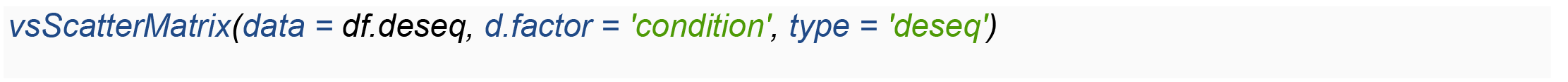

**Figure 4.**
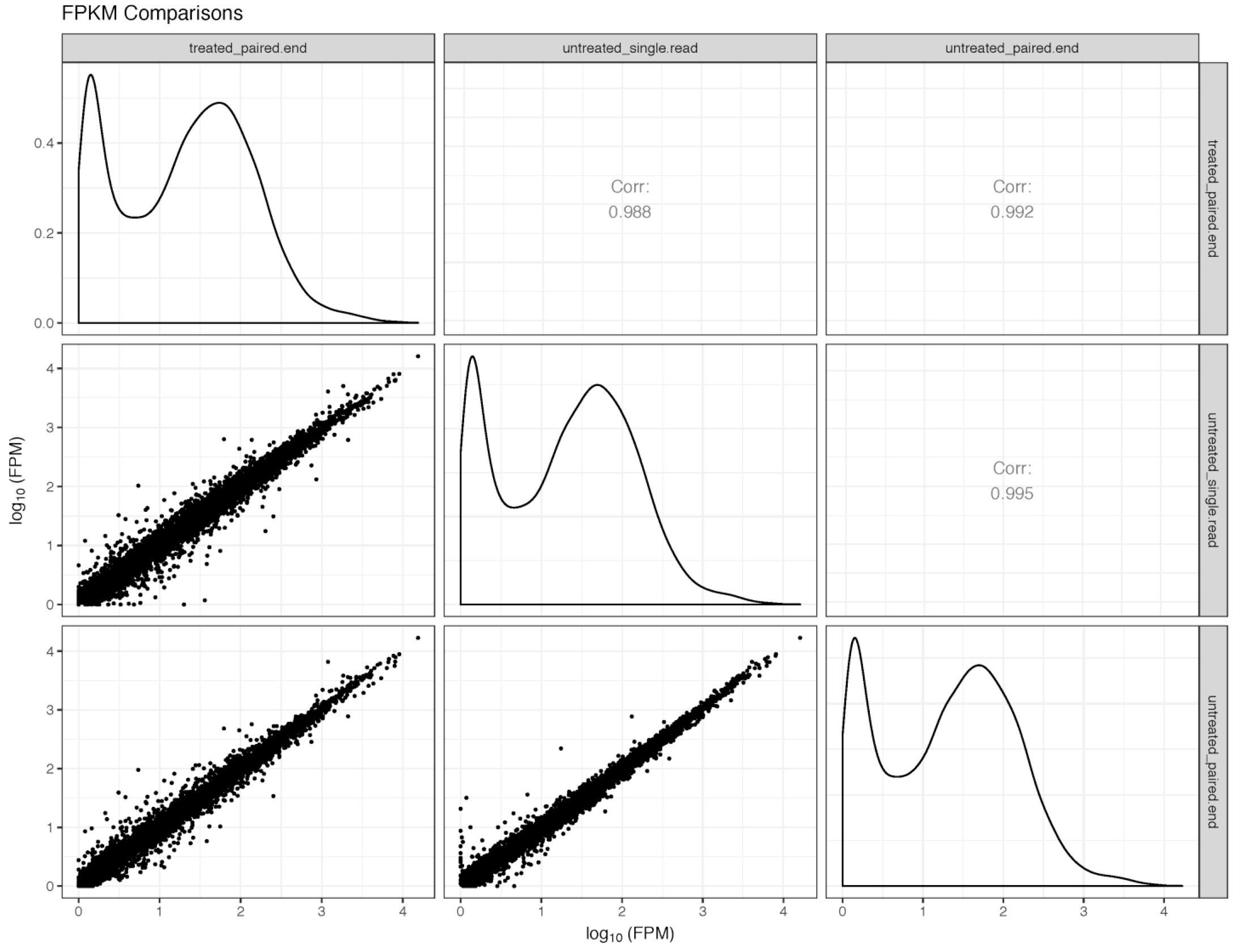
Visualization generated by the *vsScatterMatrix* function from the ViDGER package using a DESeq2 dataset, requiring a dataset, factor type, and appropriate tool type. Optional parameters include inclusion/exclusion of the main title, legend, and grid, manual specification of the main title, and manual specification of comparisons of interest. In addition to the pairwise scatterplots, density plots are provided along the diagonal and pairwise correlation values are provided in the opposite half of the matrix.

**(iv) *vsDEGMatrix*** visualizes the number of DEGs at a specified adjusted *p*-value for each treatment comparison (Figure 5, Example S5). It can be utilized to quantify the number of significantly DEGs for each comparison and provides a heatmap-based color scheme with a gradient to represent the relative magnitude of DEGs for each comparison. Like the other matrix functions, data specification and analytical type are required. The user can also specify an adjusted *p*-value which defaults to 0.05.

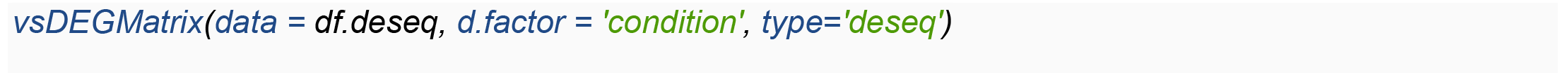

**Figure 5.**
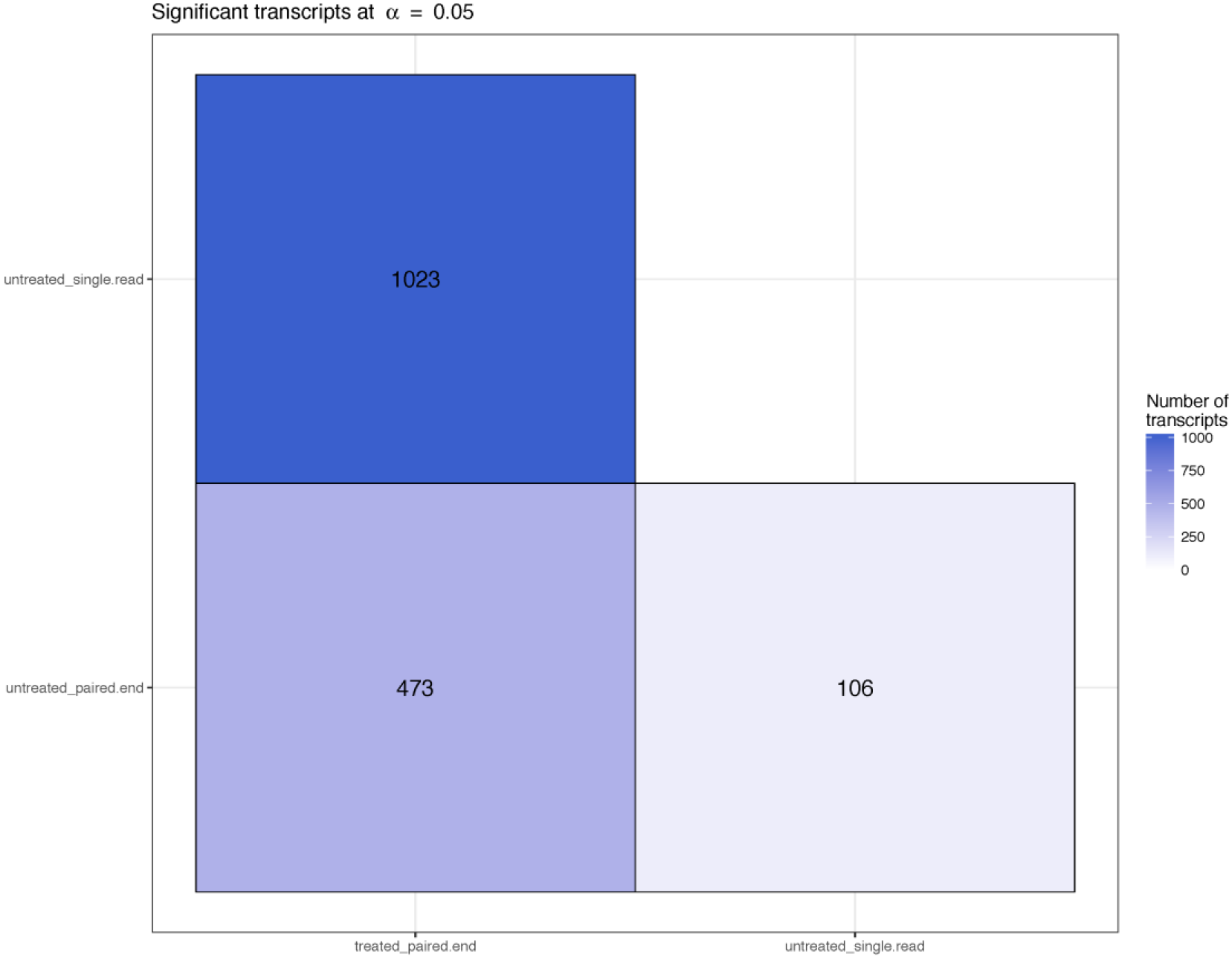
Visualization generated by the *vsDEGMatrix* function from the ViDGER package using a DESeq2 dataset, requiring a dataset, factor type, and appropriate tool type. Optional parameters include inclusion/exclusion of the main title, legend, and grid and specification of adjusted *p*-value cutoff (default is 0.05).

### Tier II Functions

**(v) *vsMAPlot*** creates an MA plot, which is a scatter plot with M (log ratio) and A (mean average) scales, of *lfc* versus normalized mean counts (Figure 6, Example S6). In addition to the basic plotting of the data points relative to the mean expression values and *lfc*, the *vsMAPlot* function also integrates visualization features that allow for a better understanding of the data. Data points in the MA plot are colored based on thresholds for the adjusted *p*-value and *lfc* of the gene in the indicated comparison to provide valuable global interpretability. Additionally, it is inevitable with most datasets that some points will be extreme relative to the majority of the data, which caused problems when generating visualizations. To address this issue, *vsMAPlot* scales the window based on the bulk of the data and represents outliers with distinct data points, indicating the magnitude of the outlier based on the size of the point. This process allows for the visualization to present the majority of the information in a viewable, usable format that is robust to outliers. Visualizing the data through this approach allows for the comparison of two treatment groups relative to the mean expression value and *lfc*. The *x* and *y* parameters specify how the fold changes are generated (e.g., *FC* = *log*_2_(sample y/sample x)).

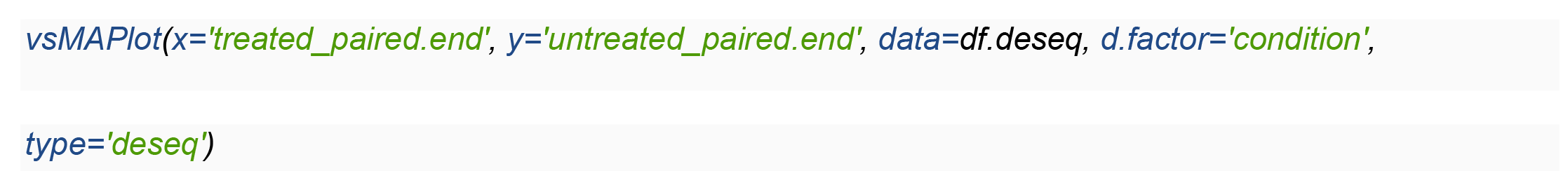

**Figure 6.**
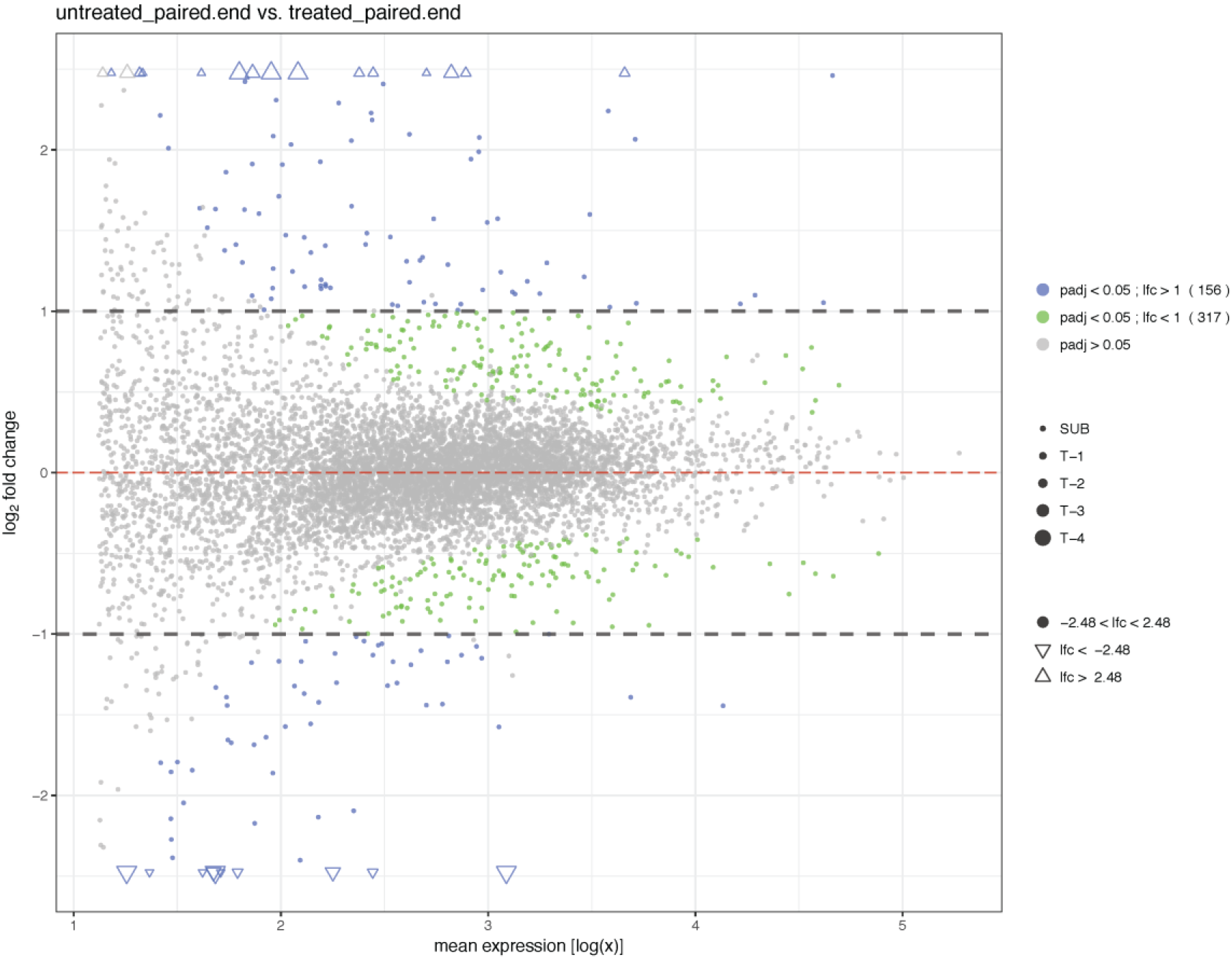
Visualization generated by the *vsMAPlot* function from the ViDGER package using a DESeq2 dataset, requiring a dataset, factor type, two factor levels, and appropriate tool type. Optional parameters include inclusion/exclusion of the main title, legend, and grid, manual specification of the y-axis limits, *lfc* threshold (default is 1), and adjusted *p*-value cutoff (default is 0.05), and specification of returning data in tabular form.

**(vi) *vsMAMatrix*** generates a matrix of MA plots for all possible pairwise treatment comparisons (Figure 7, Example S7). This process, as with the other matrix options, allows users to visualize all their treatment-based comparisons in one figure. This matrix option also includes counts for each figure based on *lfc* and adjusted *p*-value thresholds, which can be specified by the user or revert to 175 the default 1 and 0.05, respectively.

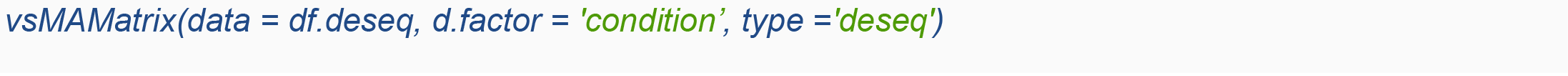

**Figure 7.**
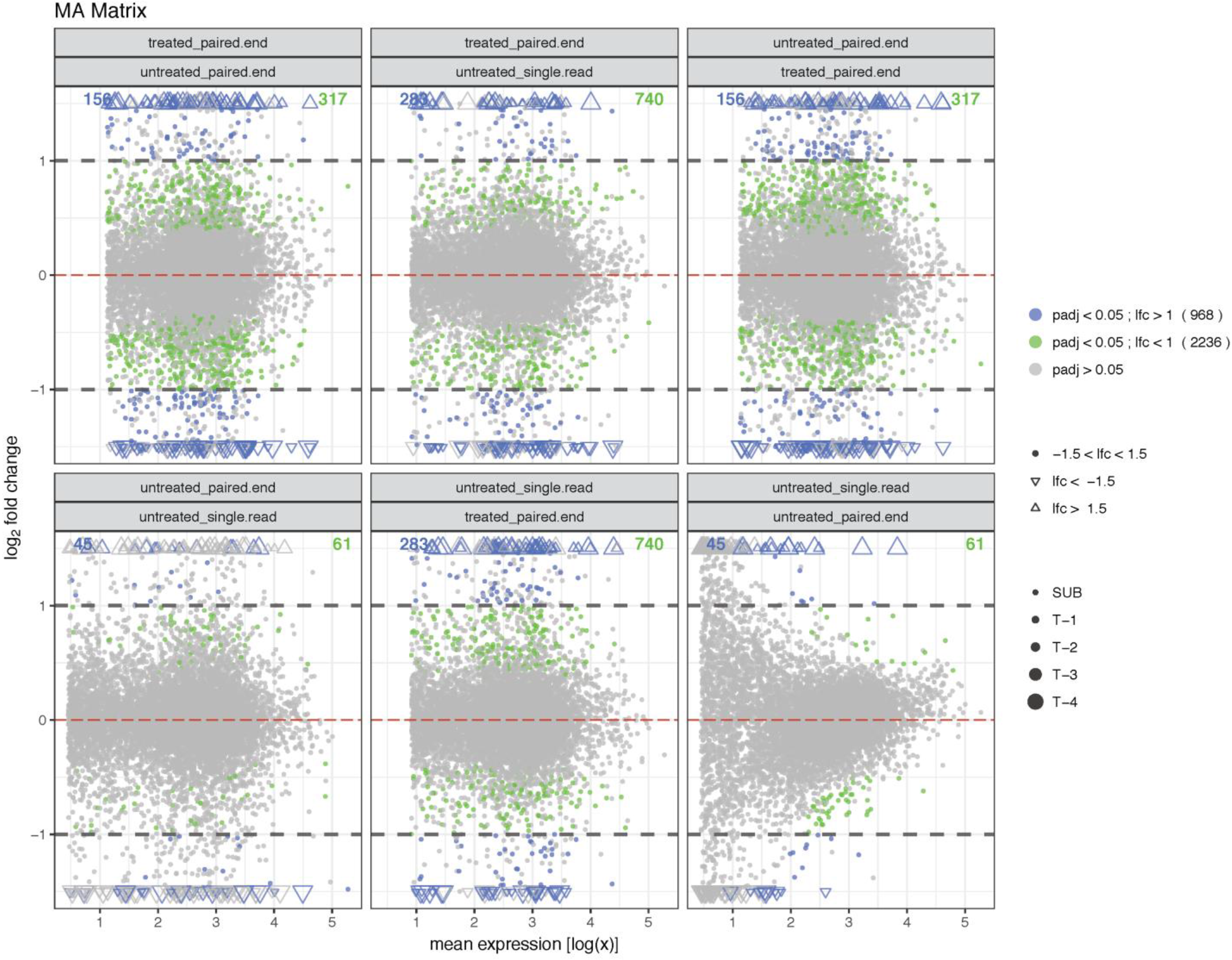
Visualization generated by the *vsMAMatrix* function from the ViDGER package using a DESeq2 dataset, requiring a dataset, factor type, and appropriate tool type. Optional parameters include inclusion/exclusion of the main title, legend, grid, and partitioned counts and manual specification of the x-axis limits, *lfc* threshold (default is 1), and adjusted *p*-value cutoff (default is 0.05).

**(vii) *vsVolcano*** creates a volcano plot for two treatments comparison by plotting the -*log*_10_(*p*-value) against the *lfc* (Figure 8, Example S8). As with the *vsMAPlot* function, the *vsVolcano* function utilizes coloring schemes to indicate the significance of magnitude of differential expression for the individual data points. Additionally, this function integrates the same data point and sizing structure to focus the plot window on the majority of the data, indicating outliers in this format.

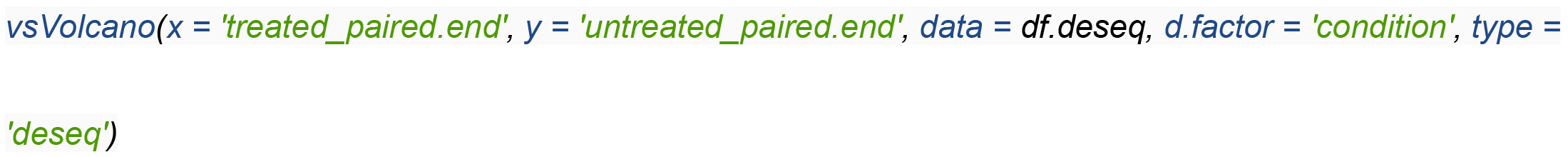

**Figure 8.**
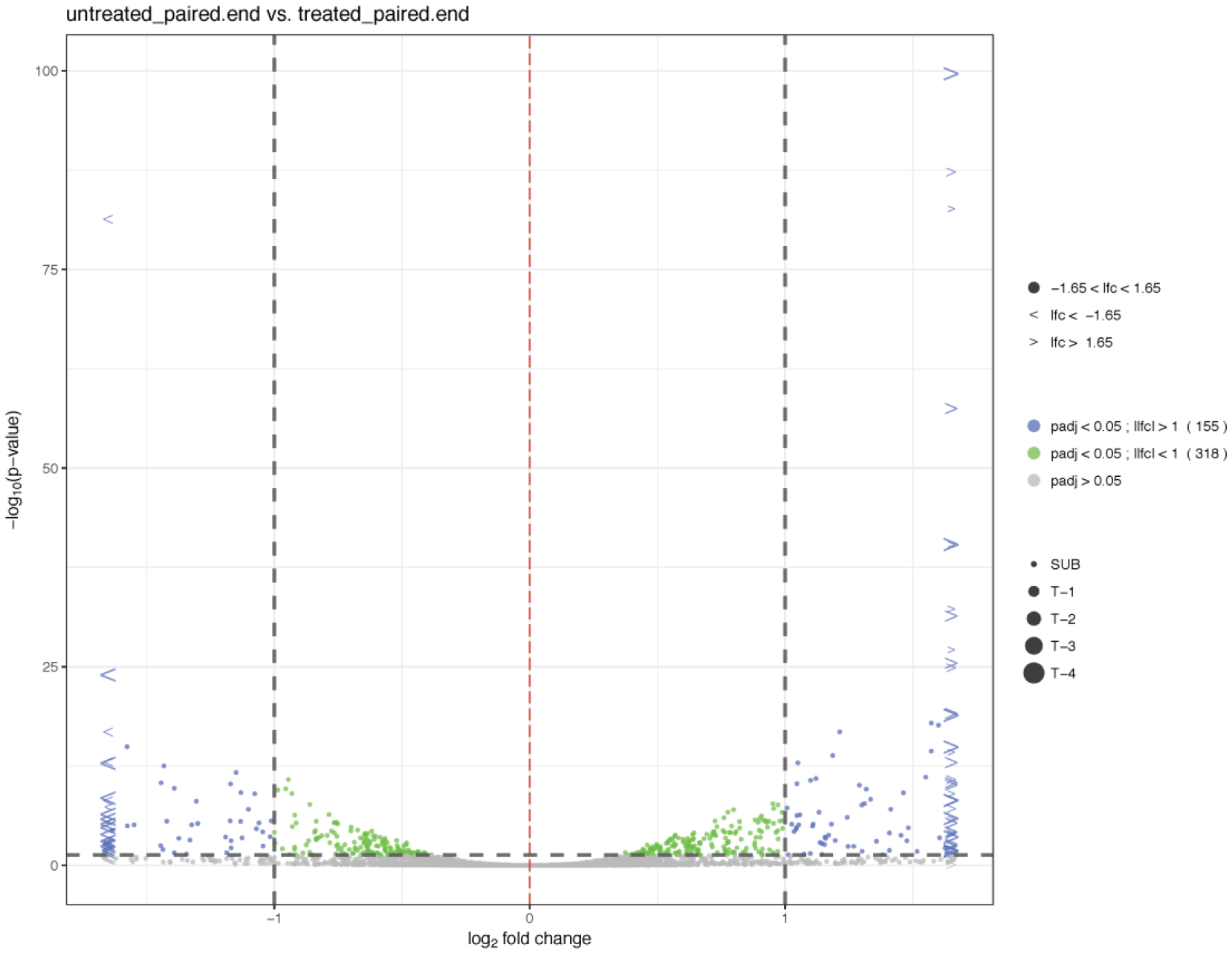
Visualization generated by the *vsVolcano* function from the ViDGER package using a DESeq2 dataset, requiring a dataset, factor type, two factor levels, and appropriate tool type. Optional parameters include inclusion/exclusion of the main title, legend, and grid, manual specification of the x-axis limits, *lfc* threshold (default is 1), and adjusted *p*-value cutoff (default is 0.05), and specification of returning data in tabular form.

**(viii) *vsVolcanoMatrix*** generates a matrix of volcano plots for all possible pairwise treatment comparison (Figure 9, Example S9). This process, as with the other matrix options, allows users to visualize all their treatment-based comparisons in one figure. Additionally, to provide a more comprehensive view with a single figure, we included a count for each separate Volcano plot based on the number of data points in each section as specified by the *lfc* and adjusted *p*-value thresholds.

Although this option may have experience limited use, it would be useful in situations where users wish to show mass similarity across all comparisons, highlight the individual or limited deviations, or display situations where the comparisons vary widely.

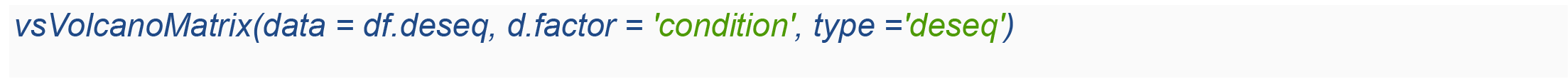

**Figure 9.**
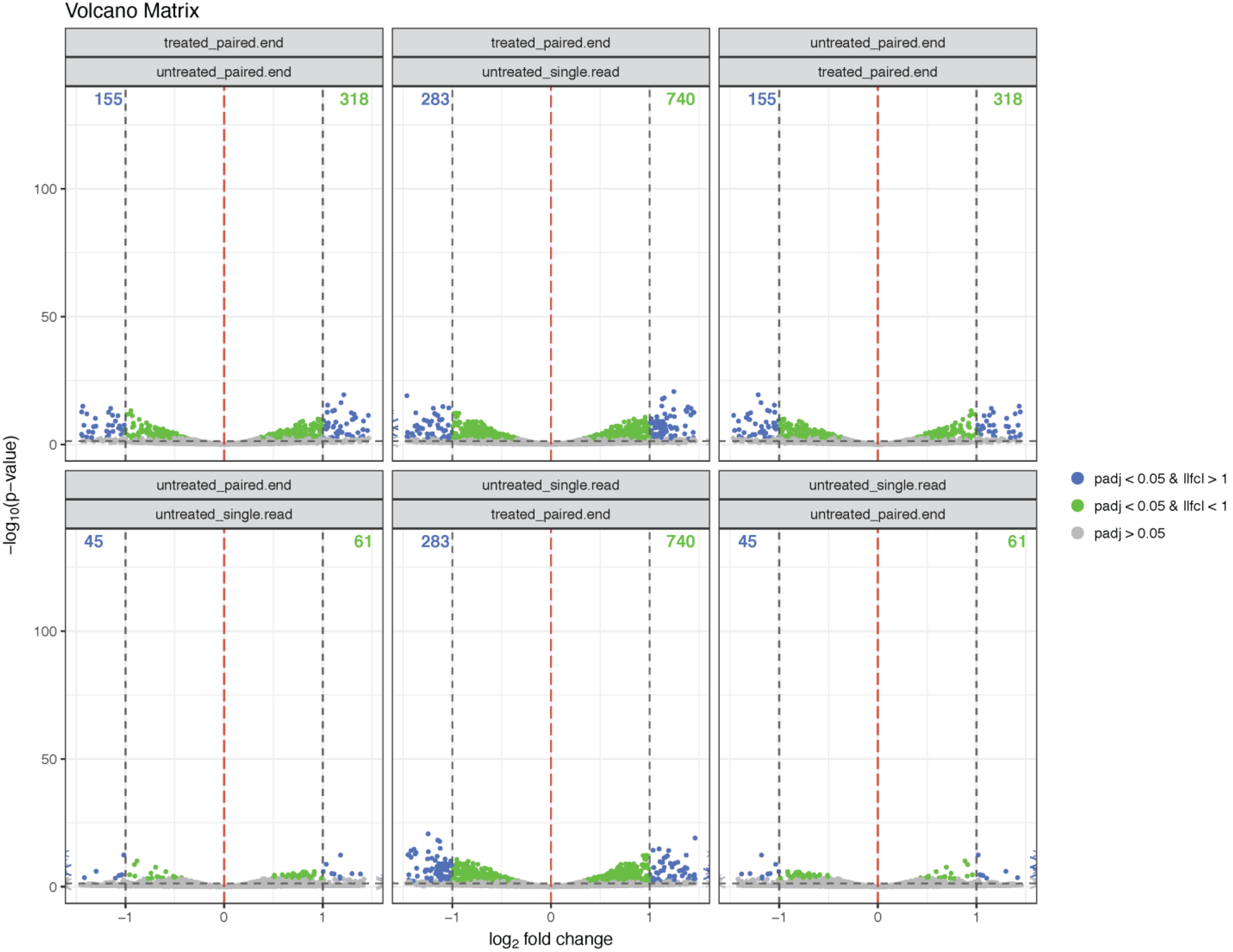
Visualization generated by the *vsVolcanoMatrix* function from the ViDGER package using a DESeq2 dataset, requiring a dataset, factor type, and appropriate tool type. Optional parameters include inclusion/exclusion of the main title, legend, grid, and partitioned counts and manual specification of the y-axis limits, *lfc* threshold (default is 1), and adjusted *p*-value cutoff (default is 0.05).

**(ix) *vsFourWay*** creates a scatter plot comparing the *lfc* between two samples and one control (Figure 10, Example S10). This approach is most useful when there are multiple comparisons being made against a specific control or relative sample. Using this function, a plot can be generated for visualizing the expression scatterplots, relative to another expression scatterplot. As with the other two main Tier 2 functions, *vsFourWay* integrates data point features to highlight significant adjusted *p*-values, over-threshold *lfc*, and outliers. In this function, *x* and *y* arguments are needed, and a *control* level is also required. Although it is possible to generate a matrix option for the FourWay plot, the authors decided against this because of two main issues. First, the *vsFourWay* function generates a significant amount of information in a single figure, with nine distinct sections representing nine distinct combinations of relative *lfc*. Creating a matrix visualization with this figure would then force each FourWay plot to be too small to collect meaningful interpretations from, thus counteracting the purpose of the package. Secondly, the *vsFourWay* function already requires three factor levels for comparison—one reference level and two comparison levels. A matrix option for this functionality would then require a minimum of four factor levels, with at least five factor levels being preferred to generate a fully-informative matrix option. This requirement would potentially put most applications out of the scope of the matrix option for the *vsFourWay* function.

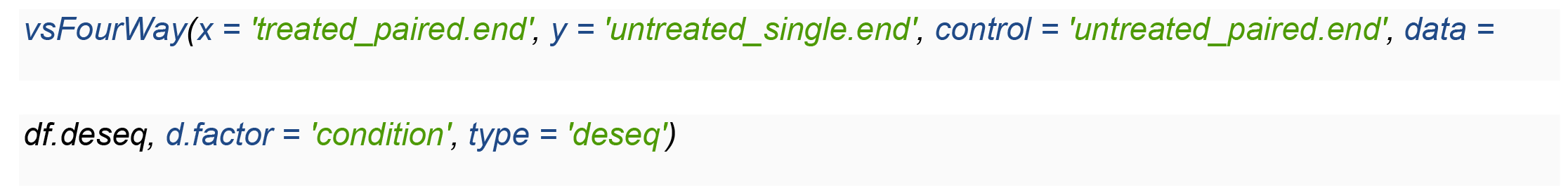

**Figure 10.**
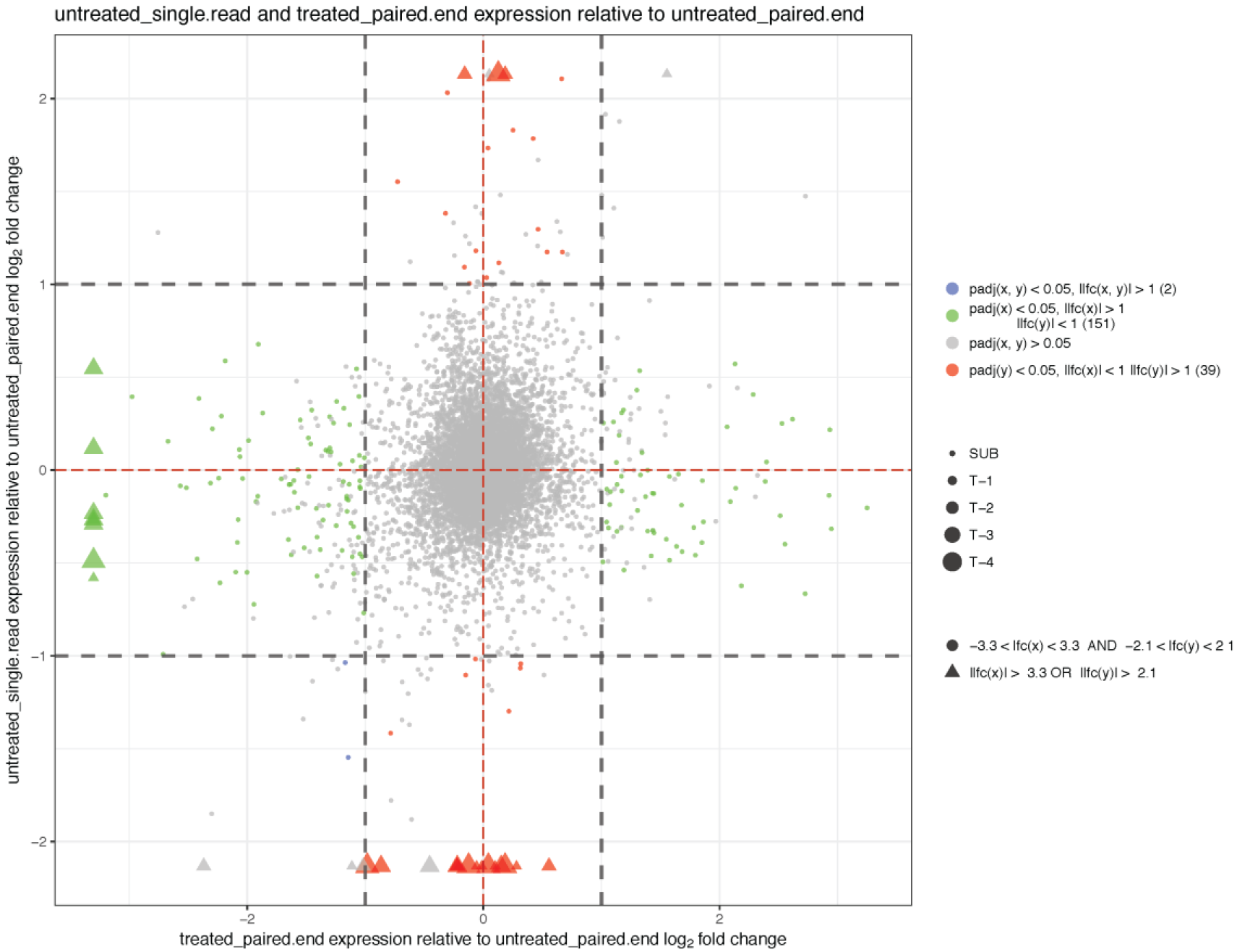
Visualization generated by the *vsFourWay* function from the ViDGER package using a DESeq2 dataset, requiring a dataset, factor type, two factor levels, reference factor level, and appropriate tool type. Optional parameters include inclusion/exclusion of the main title, legend, and grid, manual specification of the x- and y-axis limits, *lfc* threshold (default is 1), and adjusted *p*-value cutoff (default is 0.05), and specification or returning data in tabular form.

## Data Extraction

It is noteworthy that functions **(v)**, **(vii)**, and **(ix)** can return interpreting results shown in the visualizations for further analysis and interpretation (Table 2). The data extracted contains all relevant information used to generate the specified figure, including mean expression for the *x*, *y*, and *control* (in the *vsFourWay* function) factor levels, x- and y-axis values for the relevant figure, an ‘isDE’ column indicating whether the gene ID is differentially expressed based on the adjusted *p*-value threshold, ‘color’ indicating the color of the data point in the figure—which corresponds to the *lfc* and adjusted *p*-value thresholds—and ‘size’ indicating whether the data point is on the plot or an outlier and magnitude of that outlier. The data extraction is accomplished by setting the *data.return* parameter to *TRUE*.

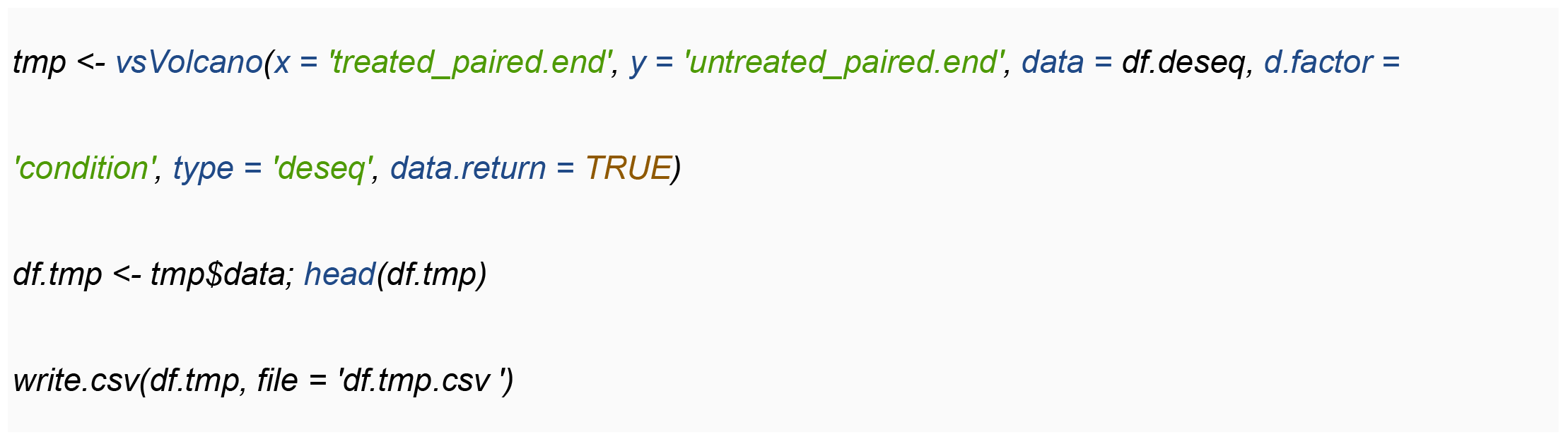
Box initially at rest on sled sliding across ice.

**Table 2.** Data extraction from the *vsVolcano* function from the ViDGER package using a DESeq2 dataset. This is the same parameterization as used in Figure 8, except *data.return* = *TRUE*. This modification will allow the user to extract relevant data from the figure. In this case, the extracted data frame includes mean expression values for the *x* and *y* factor levels, *log*_2_ fold change (logFC), *p*-value (pval), adjusted *p*-value (padj), ‘isDE’ which represents whether the differential expression is significant, ‘color’ which signifies the color of the data point corresponding to the adjusted *p*-value and *lfc* thresholds, and ‘size’ which indicates whether the data point is within the plot frame or an outlier of a particular magnitude.

|  | x | y | logFC | pval | padj | isDE | color | size |
| --- | --- | --- | --- | --- | --- | --- | --- | --- |
| FBgn0000008 | 7.922277 | 8.322253 | 0.071059 | 0.828806 | 0.974685 | FALSE | grey | sub |
| FBgn0000017 | 318.9575 | 383.2851 | 0.265054 | 0.090161 | 0.467683 | FALSE | grey | sub |
| FBgn0000018 | 30.25862 | 31.26999 | 0.047433 | 0.801233 | 0.971289 | FALSE | grey | sub |
| FBgn0000032 | 72.34193 | 72.90323 | 0.011151 | 0.949842 | 0.993072 | FALSE | grey | sub |
| FBgn0000037 | 1.539581 | 0.812298 | -0.92246 | 0.231142 | 0.700057 | FALSE | grey | sub |
| FBgn0000042 | 7928.525 | 5600.305 | -0.50155 | 0.000611 | 0.013572 | TRUE | green | sub |
| FBgn0000043 | 3273.939 | 1943.285 | -0.75253 | 7.96E-08 | 5.68E-06 | TRUE | green | sub |
| FBgn0000044 | 2.222025 | 1.599588 | -0.47417 | 0.456526 | 0.872166 | FALSE | grey | sub |
| FBgn0000046 | 2.235611 | 1.530254 | -0.5469 | 0.439278 | 0.865892 | FALSE | grey | sub |
| FBgn0000052 | 187.1546 | 201.4374 | 0.106101 | 0.498756 | 0.889058 | FALSE | grey | sub |
| FBgn0000053 | 200.4191 | 161.0824 | -0.31522 | 0.03254 | 0.260826 | FALSE | grey | sub |
| FBgn0000054 | 50.24601 | 52.8436 | 0.07272 | 0.675076 | 0.949335 | FALSE | grey | sub |
| FBgn0000057 | 56.88491 | 55.52939 | -0.03479 | 0.831612 | 0.974685 | FALSE | grey | sub |
| FBgn0000063 | 34.43974 | 27.58587 | -0.32014 | 0.084865 | 0.453512 | FALSE | grey | sub |
| FBgn0000064 | 738.3808 | 597.9759 | -0.30428 | 0.010905 | 0.125567 | FALSE | grey | sub |
| FBgn0000071 | 54.98497 | 9.358834 | -2.55464 | 1.98E-27 | 1.17E-24 | TRUE | blue | t4 |
| FBgn0000077 | 17.98973 | 17.58636 | -0.03272 | 0.898181 | 0.985072 | FALSE | grey | sub |
| FBgn0000078 | 1.743642 | 3.473474 | 0.994276 | 0.058949 | 0.37159 | FALSE | grey | sub |
| FBgn0000079 | 9.722732 | 21.87556 | 1.169886 | 3.45E-06 | 0.000156 | TRUE | blue | sub |

## Implementation

**ViDGER** is a package developed for the R environment (>= 3.3.2) and is freely available at https://github.com/btmonier/vidger. Several package dependencies are required, i.e., *ggplot2* [32], *ggally* [33], *dplyr* [34], and *tidyr* [35]. Currently, it is compatible with three commonly used DGE analysis packages, which are *Cuffdiff*, *edgeR*, and *DESeq2*. Function efficiency varies depending on what type of RNA-seq package is used. Functions used for *Cuffdiff* and *edgeR* objects complete in < 1s and while *DESeq2* objects can take up to 5s to complete. *DESeq2* objects take longer to process due to the nature of the object, which contains more stored information than the relatively simple objects for *Cuffdiff* and *edgeR*. One exception is the volcano plot matrix function **(vii)**. *Cuffdiff* and *edgeR* objects took < 10s to complete while *DESeq2* objects took >10s (Method S2). Calculations were performed on three toy data sets from *Cuffdiff*, *DESeq2*, and *edgeR* outputs. Additionally, we tested the robustness of this package on multiple large-scale RNA-seq datasets from human and plant samples (Example S1). All computations were performed on a computer with a 64-bit Windows 10 operating system, 8 GB of RAM, and an Intel Core i5-6400 processor running at 2.7 GHz.

## Conclusions

DEGs are frequently used to determine genotypical differences between two or more conditions of cells, in support of specific hypothesis-driven studies. Interpretation of this information can benefit significantly from the graphical representation of results files. We have created an R package to assist in the process of generating publication quality figures of DGE results files from *Cuffdiff*, *DESeq2*, and *edgeR*. We believe that this package will greatly assist biologists and bioinformaticians in their interpretations of DGE results. Utilizing this package will provide a straightforward method for comprehensively viewing DEGs between samples of interest and allows researchers to generate usable figures for furthered dissemination of their DGE studies.

## Key Points

- The ViDGER R package provides a straightforward method for visualizing DGE results files.
- This package integrates DGE results from the three most commonly used DGE tools: DESeq2, edgeR, & Cuffdiff.
- Nine functions are provided, including six distinct visualizations with three matrix options.
- The generated visualizations provide comprehensive views of the DGE results files in highly-informative, publication-quality figures, all of which can be extracted in multiple formats.
- ViDGER also provides a useful method for extracting relevant data from the generated figures, which is useful for further interpretation of the DGE results.

## Supplemental Materials

A supplemental file is included with this manuscript that provides more detailed information on the implementation and applications of the ViDGER R/Bioconductor package.

## Acknowledgement

We thank our collaborators for their insightful suggestions on this manuscript and pipeline testing.

